# Adaptable Fabrication of Vascularized Milliscale Tissues in Membrane-Free Organ Chips Manufactured with 3D Printed Molds

**DOI:** 10.1101/2023.12.06.570409

**Authors:** C. Ethan Byrne, K. Michael Conrad, Ashley T. Martier, Gabrielle Fortes, G. Wills Kpeli, Elisabet Olsen, William Bralower, Caroline C. Culp, Max Wendell, Keefer A. Boone, Mark J. Mondrinos

**Author notes:** These authors have contributed equally to this work (C. Ethan Byrne, K. Michael Conrad, Ashley T. Martier). Correspondence should be addressed to: Mark J. Mondrinos, Ph.D., Tulane University, 6823 St. Charles Avenue, New Orleans, LA, USA 70118. **Declarations** The research reported in this paper was funded by Mark J. Mondrinos’ faculty startup package provided by Tulane University. The authors have no conflicts of interest to report. All CAD design files, and the code used for image analyses are made available in the Supplementary Information for this article. All data and more detailed explanations of laboratory procedures will be made available upon reasonable request to the corresponding author.

## Abstract

Inexpensive stereolithography (SLA) 3D printing enables rapid prototyping of resin molds for polydimethylsiloxane (PDMS) soft lithography and organ chip fabrication, but geometric distortion and surface roughness of SLA resins can impede the development of adaptable manufacturing workflows. This study reports post-processing procedures for manufacturing SLA-printed molds built with a Formlabs F3 printer that produce fully cured, flat, patently bonded, and optically clear PDMS organ chips. User injection loading tests with iterated guide structure designs were conducted to achieve engineering reduction to practice of milliscale membrane-free organ chips (MFOC), defined as reproducible loading of aqueous solutions without failure of surface tension-based liquid patterning. The optimized manufacturing workflow was applied to further engineer milliscale MFOC for specific applications in modeling vascular physiology and pathobiology. The open lateral interfaces of bulk tissues seeded in MFOC facilitate the formation of anastomoses with internal vasculature to create milliscale perfusable vascular beds. After optimizing bulk tissue vasculogenesis in MFOC, we developed a method for seeding the bulk tissue interfaces with a confluent endothelium to stimulate self-assembly of perfusable anastomoses with the internal vasculature. Rocker- and pump-based flow-conditioning protocols were tested to engineer enhanced barrier function of the perfusable internal vasculature. Modularity of the MFOC design enabled creation of a multi-organ device that was used to model decaying gradients of cancer-associated vascular inflammation in organ compartments positioned at increasing distances from a tumor compartment. These easily adaptable methods for designing and fabricating vascularized microphysiological systems can accelerate their adoption in a diverse range of preclinical laboratory settings.

## Introduction

Early organ chips were manufactured using photolithographic methods to create silicon master molds used for downstream polydimethylsiloxane (PDMS) soft lithography and microfluidic device assembly.^1,2^ Advances in digital manufacturing now enable low-cost, rapid prototyping capabilities that eliminate the need for cleanroom access and cut the mold fabrication time from days to hours.^3^ Depending on the geometric intricacy and size scale of the design, inexpensive commercial stereolithography (SLA) based 3D printers can deliver the resolution and accuracy necessary to fabricate master molds for PDMS soft lithography.^3–5^ Lower cost and faster manufacturing of SLA printed parts comes with the tradeoffs of material property limitations. SLA resins are prone to warping upon drying and curing and have been shown to leach chemicals that can inhibit PDMS curing and injure cells cultured in fabricated PDMS culture devices.^6,7^ Furthermore, the surface roughness of SLA-printed molds negates the accurate feature replication, ease of release, and transparency of molded PDMS that are hallmarks of silicon master molds used to fabricate microfluidic devices.^8,9^ Various methods for mitigating surface roughness of 3D printed molds have been reported ^10,11^, but not within the context of a comprehensive post-processing workflow that addresses a range of material property limitations. We developed widely adaptable procedures that correct these limitations and enable efficient application-specific organ chip manufacturing in non-expert laboratories using low-cost SLA printers as a fabrication centerpiece.

The majority of reported organ chip designs can be classified as either vertically stacked layers of channels and chambers separated by porous membranes^12^, or membrane-free organ chip (MFOC) designs with horizontally arranged chambers separated by features such as pillars or guide rails that create surface tension to pattern injected hydrogels, thereby creating open and accessible tissue interfaces.^13–15^ Breaching of injected liquids beyond patterned regions established by designed guide structures is the primary MFOC failure mode.^16^ We performed systematic injection loading tests with a variety of users to quantify the effects of guide structure dimensions, bulk tissue volume, and PDMS surface properties on MFOC loading success rates. Loading success is dictated by both the designed guide dimensions and the PDMS surface hydrophobicity that influences liquid cohesion along the guide structure during injection. The rapid prototyping capabilities of SLA-based mold fabrication combined with the optimized post-processing workflow enables a design, test, and iterate approach to converge on robust designs for the downstream development of complex microphysiological models.

The ability to create bulk tissues with an anastomosed and perfusable internal vasculature make MFOC ideal venues for modeling clinically relevant aspects of vascular physiology such as barrier function, vascular inflammation, and intravascular delivery of compounds and materials.^17,18^ The optimal MFOC design identified by user testing was used as a fluidically integrated culture venue to tissue engineer milliscale bulk tissues with a perfusable internal vasculature. A two-step MFOC seeding protocol was developed in which bulk tissues are allowed to undergo vasculogenesis prior to seeding the open tissue interfaces with a confluent endothelium that facilitates autonomous formation of fluidically accessible vascular anastomoses at points of contact with the internal vasculature. We then compared unidirectional pump-driven flow and pump-free oscillatory rocking as methods of flow conditioning to enhance endothelial barrier function in perfusable bulk tissues and found that pump-driven flow is superior for rapid barrier maturation. Toward the long-term goal of developing body-on-a-chip systems, we designed a multi-organ device by connecting four MFOC modules in series using a shared fluidic channel. The multiorgan MFOC was used to model vascular inflammation in normal tissues fluidically interconnected with engineered lung adenocarcinoma tissues, a key aspect of premetastatic niche (PMN) cultivation. Collectively, we present integrated organ chip manufacturing and tissue engineering procedures that will enable researchers in a wide range of laboratory settings to create microphysiological models of vascular physiology and pathophysiology with user-defined levels of complexity.

## Results

### Milliscale Organ Chip Manufacturing Using Low-Cost SLA Printers

Resolution of the Formlabs F3 SLA printer as operated in our laboratory was determined by measuring the percent deviation from the designed dimensions of common organ chip mold features such as rectangular channels and cylindrical posts (**Supplemental Figure 1**). Positive features protruding from the primary mold surface were accurate down to 200 microns, whereas negative features sinking below the surface plane were only accurate down to the 400-to 600-micron range. This resolution range was suitable for fabrication of accurate molds in our desired milliscale size range for fluidic channels and 3D tissue chambers, but the characteristics of molds fabricated using the proprietary Formlabs resins present multiple challenges for standardizable organ chip fabrication (**Figure 1A**). Molds stored at ambient temperature prior to PDMS soft lithography retained volatiles that impeded PDMS curing (**Figure 1D**). The aspect ratio of organ chip molds exacerbated warping effects and lead to significant mold curvature (**Figure 1F**). PDMS layers produced using these molds are opaque due to surface roughness of the resin material (**Figure 1G**). We developed corrective post-processing procedures to address these complications (**Figure 1B**). These easily adaptable methods ensure complete PDMS curing, standardizable device bonding, and tunable control of PDMS optical clarity and surface hydrophobicity (**Figure 1C**). Molds were baked to remove volatiles and clamped between jeweler blocks pre-heated to 130°C to flatten the molds (**Figure 1E**). Polyurethane (PU) coating of mold surfaces corrects the downstream effects of surface roughness on PDMS clarity and hydrophobicity (**Figure 2**). Durability of master molds manufactured with this workflow was confirmed for at least 20 cycles of molding without any decrement in the quality of the resultant organ chips.

**Figure 1:**
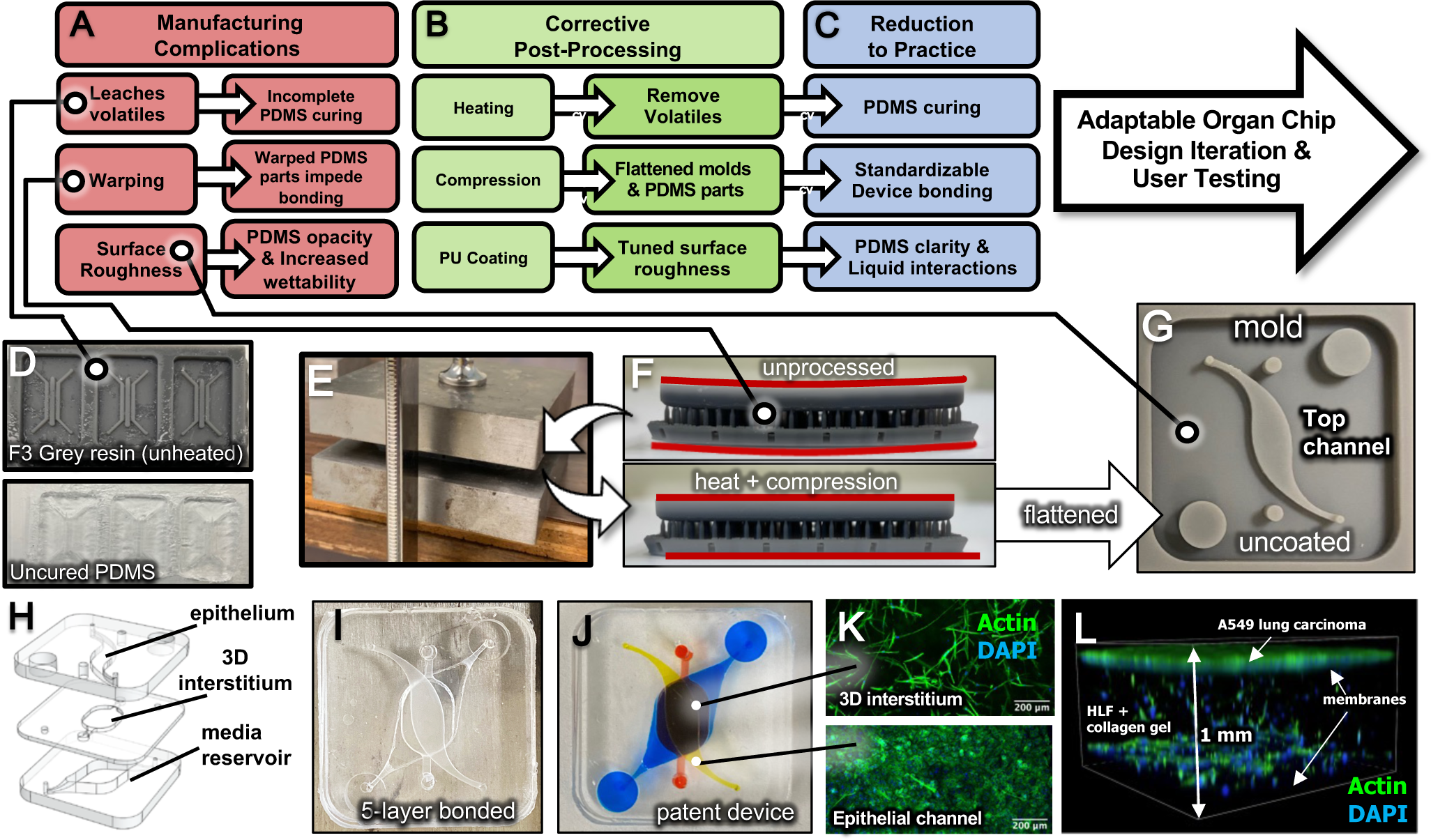
Manufacturing milliscale PDMS organ chips using a low-cost SLA printer. **A-C:** Troubleshooting and post-processing workflow for organ chip manufacturing using SLA-printed molds. **D:** Incomplete PDMS curing in SLA-printed molds with no post-processing. **E:** Clamping apparatus with heated jeweler blocks for mold flattening. **F:** Warped 3D printed mold post curing (top) compared to the same mold after baking and flattening (bottom). **G:** Flattened organ chip mold for the top layer of the multilayer device in Panels H, I and J. **H**: CAD render of exploded view of multilayer organ chip. **I:** Multilayer organ chip (3 PDMS layers, 2 membrane layers) bonded by PDMS stamping. **J:** Food dye loading to demonstrate device layering and patent bonding with flattened layers. **K:** F-actin (phalloidin, green) and DAPI (blue) labeling of the A549 cell layer and underlying 3D interstitium layer with human lung fibroblasts (HLF) embedded in collagen type I hydrogel. Scale bars = 200 μm. 7 days. **I:** 3D reconstruction (1 mm^3^ volume) of the entire 1mm height of the OIC containing A549 cells and lung fibroblasts as shown in Panel K. The position of membranes separating the device layers are indicated. Actin (green) and nuclei (blue). 7 days.

**Figure 2:**
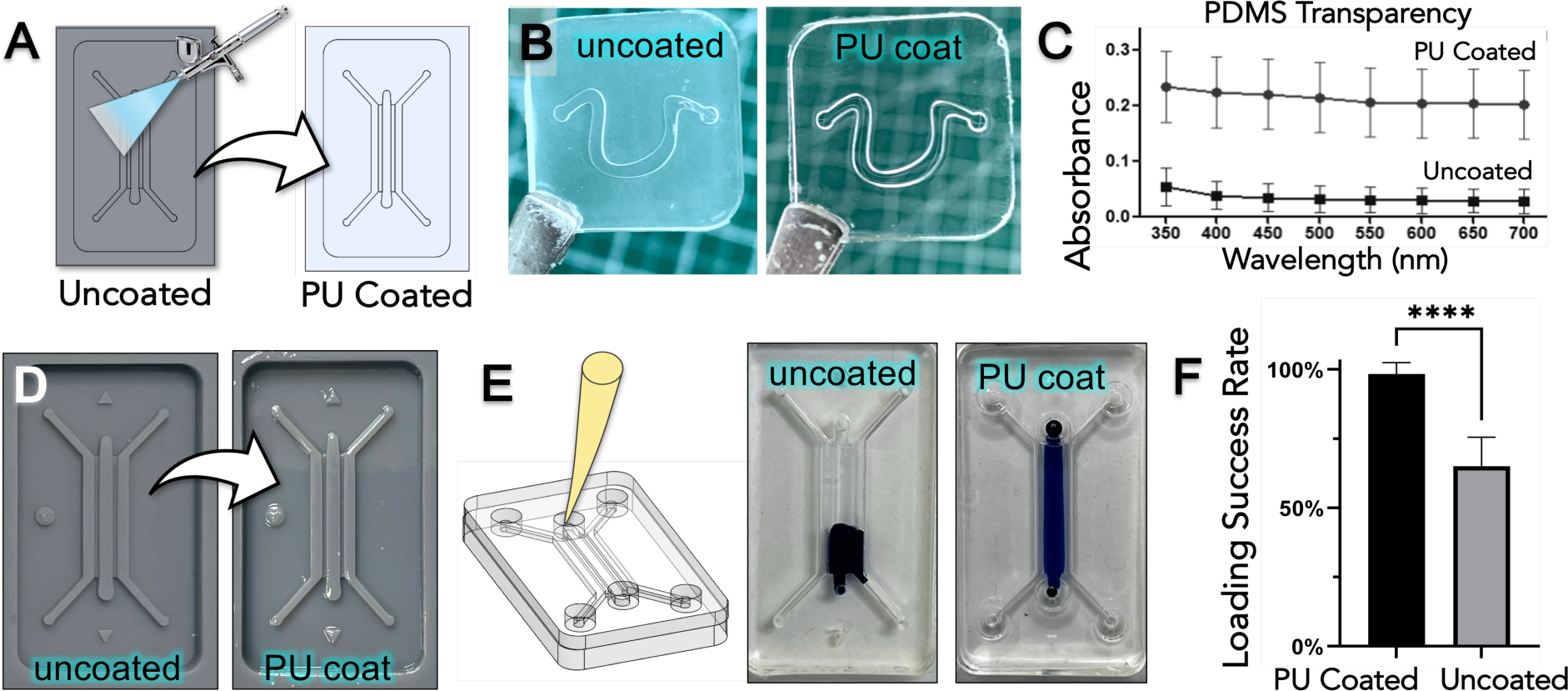
Polyurethane clear coating of SLA-printed molds to engineer PDMS clarity. **A:** SLA resin molds are polyurethane (PU) coated using a handheld airbrush. **B:** Opaque PDMS part cast in uncoated mold (left) and optically transparent PDMS cast in clear coated mold (right). **C.** Absorbance of light at varying wavelengths between clear coated and uncoated molds demonstrates the optical transparency of PDMS layers from PU coated molds. **D:** Uncoated (left) and PU-coated (right) membrane-free organ chip (MFOC) molds. **E:** Example of failed injection defined as liquid breaching over guides in device made of uncoated molds (left) and an example of successful injection in a device manufactured using coated molds (right). **F:** Comparison of success rate of loading between devices made using clear coated versus uncoated molds; n = 6 users, 10 trials each. **** indicates P < 0.0001.

To test the ability of this manufacturing workflow to produce functional organ chip models, we engineered a 5-layer organ chip (3 PDMS layers, 2 polyester membranes) for modeling the interface of an epithelium and the underlying interstitium using as-printed rough surface molds (**Figure 1H**). Patent devices were reproducibly fabricated using stamped polyester membranes for layer partitioning and did not show evidence of leakage between layers due to bonding imperfections (**Figure 1I, 1J**). We used this device to create a prototype model of the interface between lung carcinoma and an adjacent lung interstitium by seeding A549 lung adenocarcinoma cells in the upper channel and normal human lung fibroblasts embedded in type I collagen hydrogel in the 3D tissue chamber (**Figure 1K**). Uniformly robust adhesion and growth of both cell types was observed throughout the more than 1 mm vertical height of the patterned tissue interface (**Figure 1L, Supplemental Video 1**).

Many organ chip applications rely on imaging that requires optical clarity of the PDMS devices. Optical clarity is inherent to the smooth PDMS surfaces generated using standard silicon master molds. PDMS opacity resulting from the rough surfaces of as-printed SLA resin molds was mitigated by clear coating the molds with polyurethane (PU) used for automotive detailing to produce a smooth surface finish (**Figure 2A and 1D**). This manufacturing workaround yields optically clear PDMS parts (**Figure 2B**). Spectrophotometry quantitatively verified the significantly increased transparency of PDMS molded from PU-coated resin mold surfaces (**Figure 2C**). PDMS surface roughness also influences hydrophobicity and therefore liquid interactions. While liquid patterning in membrane-free organ chips (MFOC) is accomplished via surface tension created by the guide structures^14^, the PDMS surface hydrophobicity influences cohesion and therefore stability of liquid pinning along the guide structures during injection. We tested the hypothesis that loading efficiency would be decreased in MFOC with rough PDMS surfaces from uncoated molds, based on the notion that the increased hydrophobicity will tend to repel injected liquids, thereby counteracting the cohesion needed for stable pinning along the guide structure.^19^ Increased surface roughness and hydrophobicity of PDMS surfaces from uncoated molds were confirmed using DIC imaging and contact angle measurements (**Supplemental Figure 2**). Scanning electron microscopy (SEM) revealed a markedly smoother surface texture of PU-coated molds compared with uncoated molds, but microscale surface fracturing was observed on the PDMS parts from both PU coated and uncoated molds (**Supplemental Figure 3**). We quantified loading success rates of MFOC produced with uncoated or PU-coated molds (**Figure 2D and 2E**). Injection of rough-surfaced devices from uncoated molds resulted in more frequent injection loading failures defined by breaching of liquid over internal guide structures (**Figure 2F**). PU coating of molds yielded MFOC with 98 +/− 4% loading success rates compared to 65.0 +/− 11% for rough surfaced MFOC (**Figure 2G**).

### Membrane-Free Organ Chip Reduction to Practice

We aimed to engineer milliscale MFOC that enable reproducibly successful injection loading by systematically varying geometric design features and performing user testing (**Figure 3 and Supplemental Video 3**). Previously reported microscale MFOC using continuous horizontal guide structures termed ‘phase guides’ were designed with the guide structure height at 25% the height of the bulk tissue chamber (H), creating an open tissue interface height (h) that is 75% of the bulk tissue height (h/H = 0.75).^15,20^ We found that h/H = 0.75 is not tractable on the millimeter scale in pilot studies based on complete failure to pin liquids in the bulk tissue chamber. We used h/H = 0.5 as a starting point for milliscale design iteration and compared single guide and double guide designs (**Figure 3B and 3C**). Loading success rates of 85-90% were achieved in both designs by non-expert users (**Figure 3D**). The double guide configuration was chosen for further design iteration and testing due to the advantage of a symmetrically positioned central interface with all z-direction layers of the bulk tissue situated within 0.25 mm of the open bulk tissue interface.

**Figure 3:**
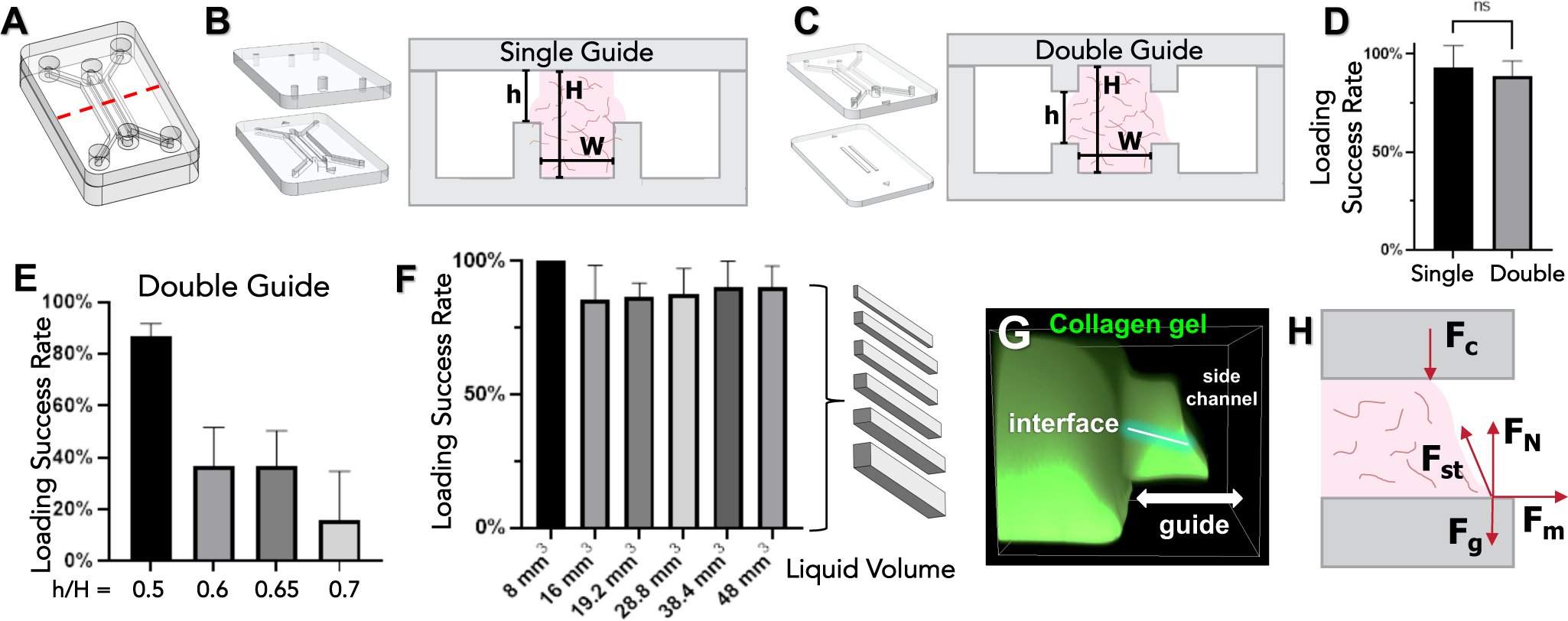
MFOC design iteration and reduction to practice through user testing. All testing consisted of least n = 6 users performing 10 injection trials per design. **A:** CAD render of a prototypical MFOC device. Dimensions of internal features and volumes were varied for user testing of loading success rate. **B&C:** CAD render of layers and drawing of cross section of single guide MFOC (**B**) and double guide MFOC (**C**). Key dimensions include guide height (h), tissue chamber height (H), and tissue width (W). A full listing of dimensions for all designs tested in this study are available in the **Supplementary Information** for this article. General size ranges for single guide MFOC: H = 0.5 mm, l = 12 mm, and W = 1-2 mm; for double guide MFOC: H = 1 mm, l = 12 mm, W = 1.2 mm. Guide width = 0.55 mm for all designs. **D:** Loading success rates for single guide and double guide MFOC. **E:** Loading success rates for various h/H values in double guide MFOC. **F:** Loading success rates for tissue volumes ranging from 8 to 48 mm^3^. H and W were varied with constant l = 12 mm. **G:** Acellular type I collagen hydrogel containing 40 kDa for visualization FITC-after injection loading and polymerization in an MFOC. Indicated guide width = 0.55 mm. **H:** Proposed free body diagram of forces acting on a liquid (collagen precursor solution for our applications) pinned between the guide structures of a double guide MFOC as shown in Panel F. F_c_ is the cohesion of the liquid and PDMS surface acting at all points of contact. F_N_ is the normal force acting at all surface contact points. F_st_ represents the surface tension forces stabilizing the liquid face. F_g_ is gravity and F_m_ is the inertial force tending to push the mass of liquid over the guide during injection loading.

Loading efficiency dropped below 40% with h/H = 0.6 or 0.65 and dropped below 20% with h/H = 0.7 (**Figure 3E**). Thus, we used h/H = 0.5 as a MFOC design specification and tested the effect of increasing the tissue chamber volume by increasing the values of H and chamber width, W. Increased inertial forces derived from a greater mass of liquid being injected would be expected to cause breaching of injected liquids over the guide structures during injection (F_m_ in **Figure 3H**). We found that increasing the tissue volume up to 6-fold did not significantly reduce the loading efficiency (**Figure 3F**). FITC-dextran loaded collagen hydrogel was cast in the base 8 mm^3^ design to visualize the formation of a foot along the bottom guide that consistently creates a downward sloping geometry at the tissue boundary (**Figure 3G**). Based on these collective observations from MFOC loading tests, we propose that stable liquid patterning requires a balance between the inertial force generated by the momentum of the injected liquid tending to drive liquid over the guide and the combined effects of the liquid surface tension force and the cohesion force between the liquid and PDMS surface (F_m_, F_st_, F_c_ in **Figure 3H**).

### Engineering Milliscale Tissues with Perfusable Bulk Vasculature in MFOC

Engineering conditions that promote the formation of a uniformly distributed 3D vascular network within tissues seeded in MFOC is the first step to fabricating a fluidically integrated bulk vasculature. We used a combination of human umbilical vein endothelial cells (HUVEC) and human lung fibroblasts (HLF) in a collagen type I and fibrin hydrogel blend to facilitate bulk tissue vasculogenesis in MFOC (**Figure 4A and 4B**). 3D vascular network assembly was evident by 3 days in culture using this formulation (**Figure 4C**). Interconnected networks spanning the entire tissue bulk assemble by 7 days in culture (**Figure 4D**). Evidence of tissue maturation was seen in the formation of perivascular niches with closely associated and enrobing fibroblasts (**Figure 4F**), and the formation of continuous laminin-rich basal lamina enrobing the vascular network (**Figure 4G**). Morphometric evidence of maturation included decreasing non-participating endothelial cell numbers, increasing vessel diameters, and increasing diffusion distances (network spacing) (**Figure 4E and 4H**).

**Figure 4:**
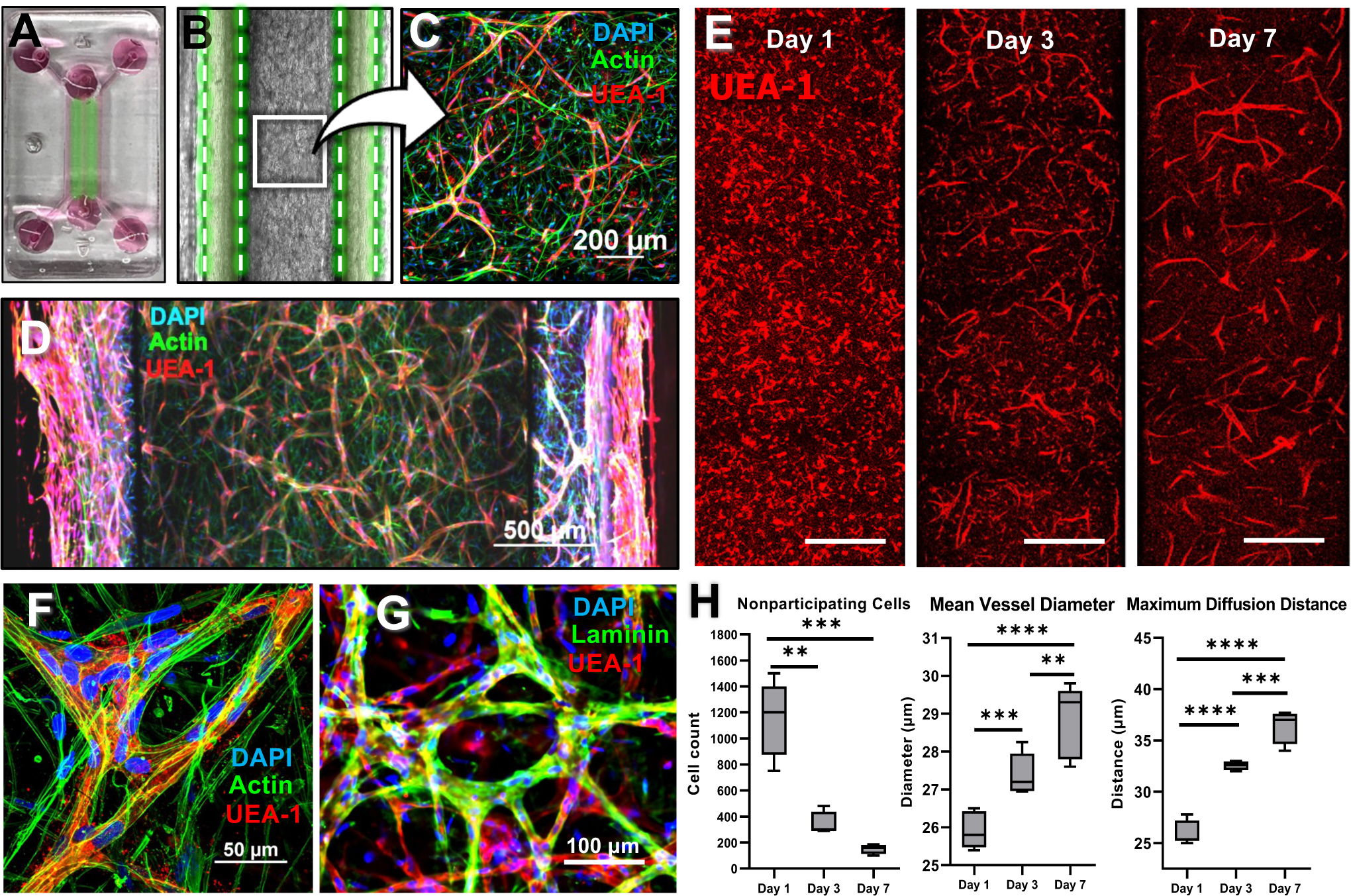
Engineering bulk tissue vasculogenesis in milliscale MFOC. **A:** Digital photograph of a MFOC loaded with cell-laden hydrogel and culture medium. Green lines indicate the position of guide structures that confine the bulk tissue. **B:** Differential interference contrast (DIC) micrograph of an MFOC loaded with HUVEC and HLF in a blend of collagen and fibrin hydrogel as described in **Materials and Methods**. Guide regions are marked by green coloration between the dashed lines. The bulk tissue is visible between the guide structures. **C:** 3D laser scanning confocal microscopy (LSCM) stack of a nascent vascular network in the bulk tissue after 3 days of culture. Actin in all cells is labeled with phalloidins (green) and endothelial cells are specifically labeled with UEA-1 lectin (red). Fibroblasts are green only. Scale bar = 200 μm. **D:** Formation of a continuous vascular network throughout the bulk tissue after 7 days of culture. Scale bar = 500 μm. **E:** Representative stitches of UEA-1 staining in 3D LSCM stacks along the bulk tissue length used for image analyses shown in Panel H. Scale bars = 500 μm. **F**: Perivascular niche occupied by fibroblasts enrobing endothelial tubules after 7 days of culture. Scale bar = 50 μm. **G:** Vascular basement membrane formation after 7 days of culture. Laminin (green), UEA-1 lectin (red), DAPI (blue). Scale bar = 100 μm. **H:** Morphometric analysis of bulk tissue vasculogenesis at Days 1, 3, and 7. The number of non-participating endothelial cells decreases with the formation of an interconnected vascular network. Mean vessel diameter increases with cell recruitment and lumen formation in the network. Max diffusion distance increases with cell coalescence into the network and network pruning.

Achieving anastomosis of the internal bulk vasculature with inlets and outlets to liquid medium in the MFOC side channels is the critical step to achieve tissue perfusion. Surfaces of the bulk tissue interfaces must be endothelialized to establish a complete vascular barrier that prevents free diffusive transport between the liquid medium and the bulk tissue volume. Side channel seeding with a high density of endothelial cells (at least 8 x 10^6^ cells/ml, see **Materials and Methods)** after 2 days of bulk tissue vasculogenesis results in immediate and complete coverage of the tissue interfaces (**Figure 5B**). Numerous anastomoses with the internal vasculature form de novo by 9 days in culture (**Figure 5C**). 3D reconstructions revealed that anastomoses appear as open pipe faces on a manifold along the length of the tissue boundary upon 3D visualization (**Figure 5C** inset). Internal vessels are inaccessible in devices without the side channels seeded, due to the formation of dense cell layers aligned parallel to the tissue boundary (**Figures 5D and 5E**).

**Figure 5.**
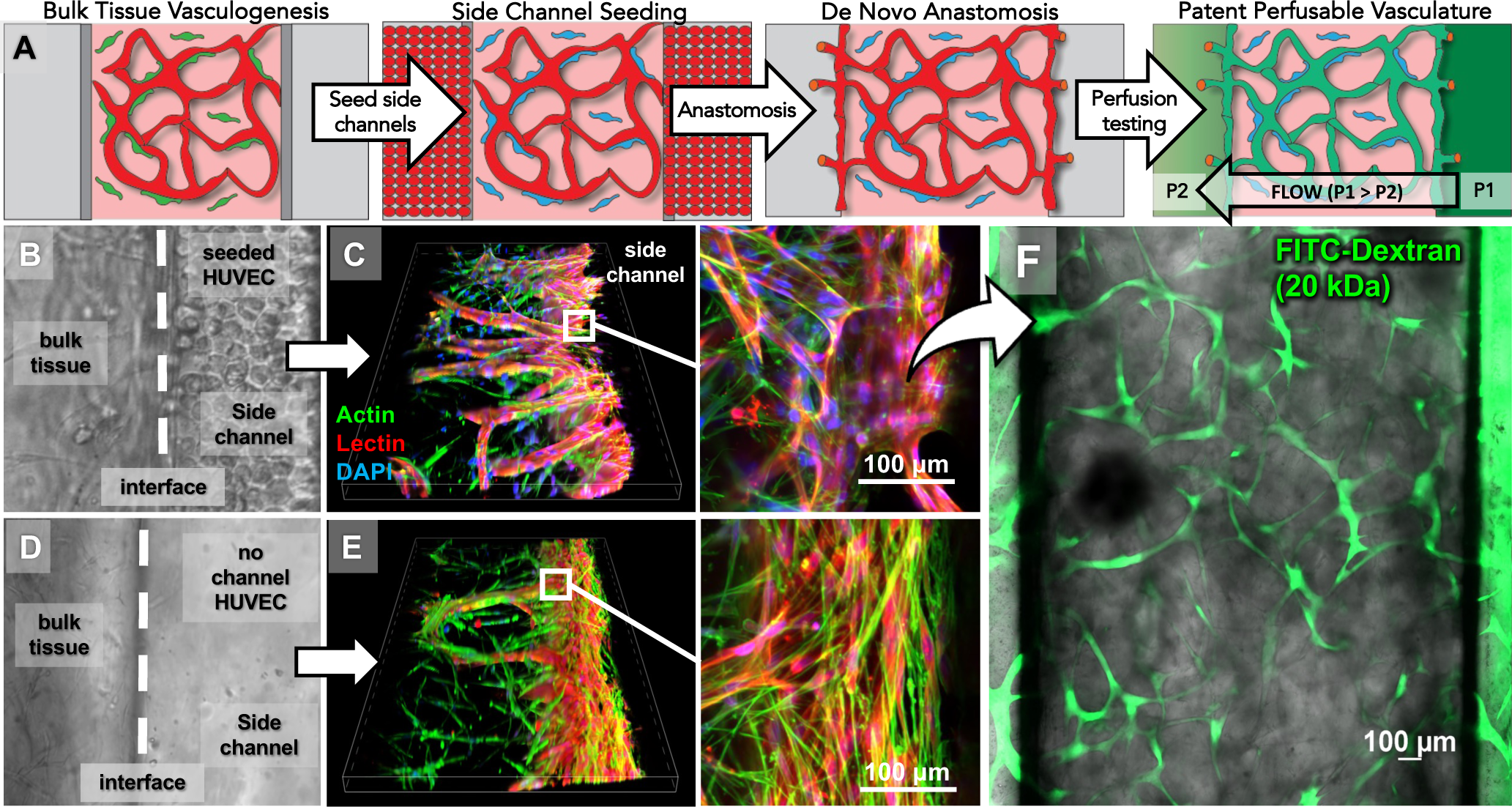
Fabricating milliscale bulk tissues with an anastomosed internal vasculature in MFOC. **A:** Workflow for establishing patent and perfusable vasculature tested by FITC-dextran perfusion as described in **Materials and Methods. B:** Differential interference contrast (DIC) image taken at the tissue interface situated on the guide structure immediately after side channel seeding (Step 2 in Panel A, after 48h of vasculogenesis). **C:** 3D LSCM stacks depicting points of vascular anastomosis at the tissue interface after 9 total days of culture. Endothelial cells are labeled with UEA-1 lectin (red). Actin in all cells is labeled with phalloidins (green). **Inset:** Open face of a single anastomosed vessel. Scale bar = 100 μm. **D:** DIC image taken at the tissue interface situated on the guide structure in a device without side channel seeding after 48 hours. **E:** 3D LCSM stack of the tissue interface in a device without side channel seeding after 48 hours. **Inset:** Dense layers of cells cover the interface and internal vascular structures remain inaccessible. 9 days of culture. Scale bar = 100 μm. **F:** Perfusion testing with 40 kDa FITC-dextran after 9 days. LCSM stack depicting a snapshot after 5 minutes of tissue perfusion. Green fluorescence is contained within vessels and absent in the interstitial spaces. Scale bar = 100 μm.

The anastomosed vasculature was perfusable and patent after 9 days of static culture (7 days after side channel seeding) as demonstrated by uniform perfusion of FITC-conjugated dextran (**Figure 5F**). While establishment of perfusable vasculature without gross leakage was attainable using this protocol, it is well-established that the fluid shear stress from flow positively regulates endothelial barrier function.^21^ We compared unidirectional syringe pump-driven flow with the more convenient rocking method that produces oscillatory bidirectional flow but does not require external connections with the device (**Figure 6**). Syringe pump-driven flow pulling at a rate of 20 μl/hour across the bulk tissue for 96 hours generated a patent vasculature as verified by perfusion testing (**Figure 6A**). While the position of anastomoses is not patterned in guide rail based MFOC, anastomoses form in a regular pattern along the 10-mm length of the device (**Figure 6C**). Perfusion of finer vessels is visible at higher magnification with longer exposure times (**Figure 6C** inset). Rocking for 96 hours yielded inconsistent results, ranging from lack of perfusion to perfusion with gross evidence of leakage into the interstitial spaces (**Figure 6B**). Perfusion in pumping conditioned devices demonstrated subjectively less leakage into the extravascular space.

**Figure 6:**
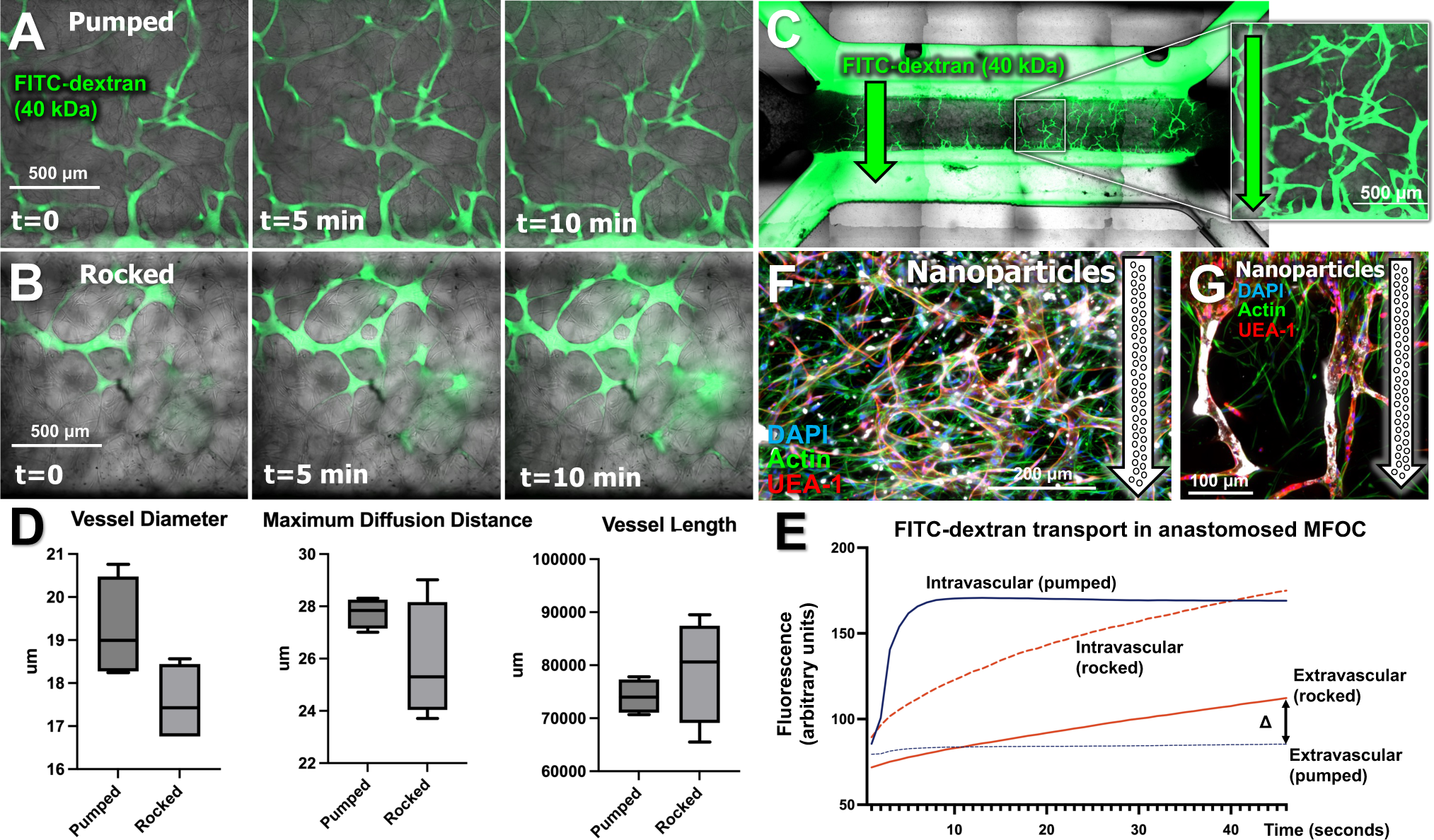
Fluid flow conditioning of anastomosed vasculature in MFOC. **A:** Time lapse images (0, 5, 10 minutes) of perfusion with 40 kDa FITC-dextran after 96 hours of syringe pump-driven flow across the bulk tissue at a rate of 20 μl/hour. Scale bar = 500 microns. **B:** Time lapse images (0, 5, 10 minutes) of perfusion with 40 kDa FITC-dextran after 96 hours of oscillatory rocking at an angle of 15 degrees and a rate of 1 cycle/minute. Scale bar = 500 microns. **C:** Composite stitched image after 10 minutes of FITC-dextran perfusion in a pumped device with patent vascularization that establishes a fluidic connection between the side channels. That communicates between liquid channels. **Inset:** A region of interest imaged at higher magnification with constrast enhancement reveals perfusion of finer vessels not visible in low magnification composites. Scale bar = 500 microns. **D:** Morphometric analysis of FITC-dextran perfused vessels in pumped and rocked devices did not reveal significant differences. **E:** Mean fluorescence in the intravascular and extravascular spaces of LCSM stacks were calaculated to assess permeability of the perfused vasculature. Recruitment of vessels for perfusion occurs more rapidly in pumped devices compared to rocked devices and there is less diffusion into the extravascular space. Data shown from a representative perfusion under each condition. Similar results were obtained in at least 3 independent replicates. **F:** Cy5-labeled lignin-g-PLGA nanoparticles accumulated in both the intravascular and extravascular spaces. Scale bars = 200 microns.

Analysis of network permeability from time course imaging revealed rapid recruitment and establishment of maximum intravascular fluorescence in perfused vessels of pumped devices upon introduction of FITC-dextran, whereas the extravascular fluorescence in pumped devices increased only slightly during the same time increment (**Figure 6E, Supplemental Video 4**). In contrast, vessel recruitment was delayed in rocked devices, and extracellular fluorescence gradually increased during the time course of perfusion, indicating vascular leakage due to incomplete endothelial barrier formation (**Figure 6E**). There were subjective differences in the morphology of pumped vs. rocked vasculature, with measured trends toward increased vessel diameter (greater perfusion volume, less resistance to flow), increased diffusion distances (greater intersegment spacing), and shorter vessel length (more pruned networks) in the vasculature of pumped devices (**Figure 6D**). The flow-conditioned vasculature in pumped MFOC was then tested as a conduit for delivering nanoparticles to the 3D bulk tissue. Nanoparticles were distributed throughout the bulk vasculature, with larger aggregates wedged near branchpoints within the vasculature and smaller particle aggregates present in the extravascular spaces (**Figure 6F**). A layer of adherent particles persists on large vessels near anastomosis points following the perfusion (**Figure 6G**).

### Design and Fabrication of Multi-Organ MFOC for Modeling Premetastatic Niche Cultivation

Toward the long-term goal of developing vascularized body-on-a-chip systems, the 8 mm^3^ single MFOC design was used as the basis for engineering a prototype multiorgan model. A device containing four single MFOC modules arranged in series with a shared fluidic communication channel was designed to create a prototype model of lung cancer (LC) premetastatic niche (PMN) cultivation that connects an engineered LC tumor microenvironment with three downstream tissue niches (TME and PMN-1,2,3 in **Figure 7A**). Control devices replaced the TME with a fourth PMN. Vascular inflammation is a key component of PMN cultivation that enables transport of serum proteins and trafficking of immune cells to the nascent PMN.^22,23^ Intense ICAM-1 staining was observed throughout the vasculature, tumor cells, and fibroblasts in the primary TME, where the concentration of tumor-derived factors is highest (**Figure 7B, TME**). ICAM-1 signal intensity in micrographs decreased qualitatively with increasing diffusion distance from the TME (**Figure 7B, PMN-1,2, and 3**). We first pooled the 3 PMN for global analysis of vascular inflammation in devices containing a TME and measured a significant increase in total ICAM-1 fluorescence intensity masked to UEA-1 lectin-labeled endothelial cells (**Figure 7C**). A linear gradient of decreasing ICAM-1 signal intensity masked to lectin-labeled endothelial cells from the TME to PMN-3 was observable and quantifiable (**Figure 7D**). A similar gradient of decaying relative non-vascular inflammation was measured by excluding the contribution of lectin-masked ICAM-1 signal (**Figure 7E**).

**Figure 7:**
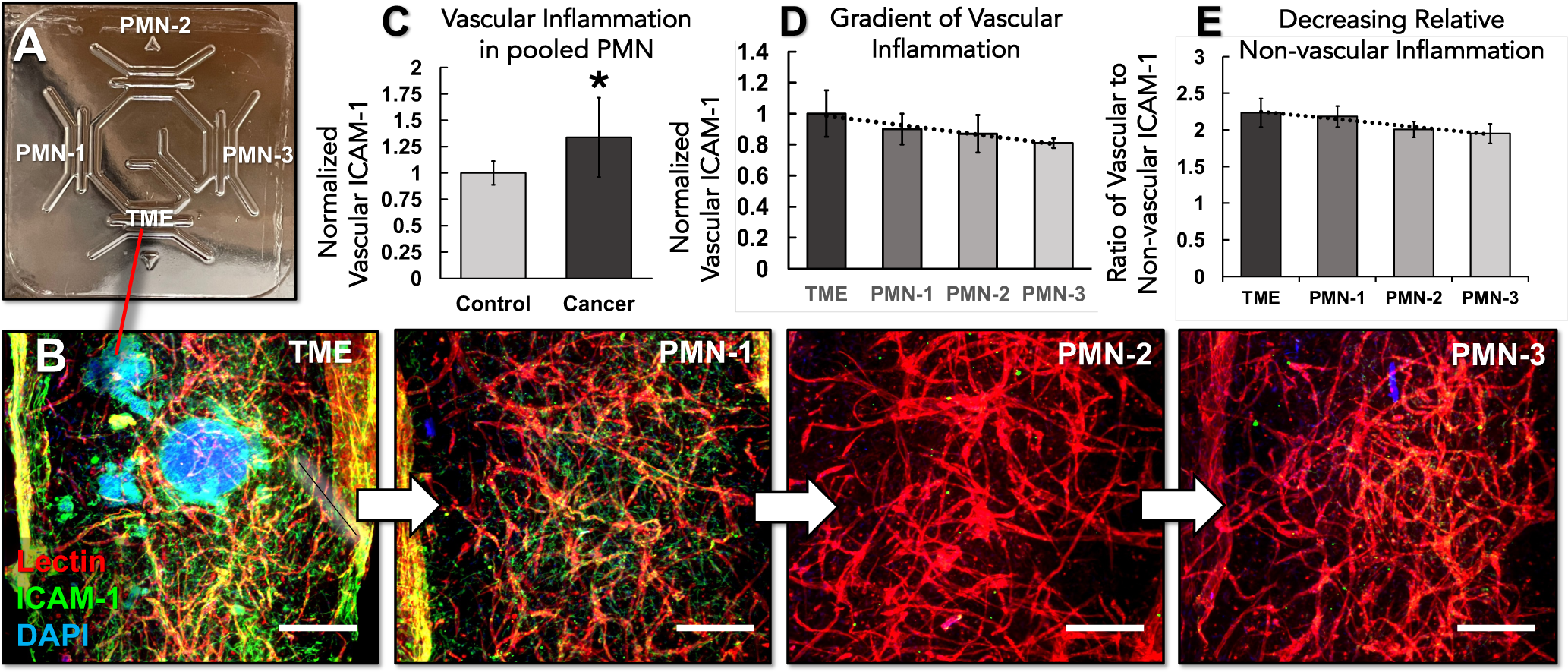
Engineering a multi-tissue MFOC for modeling of lung cancer premetastatic niche (LC-PMN) cultivation. **A:** Digital photograph of the LC-PMN device. See **Supplemental Information** for design specifications. Compartment labels indicate locations of tissues depicted in Panel B. TME = tumor microenvironment. PMN = premetastatic niche. **B:** Inflammatory activation visualized in 3D LSCM stacks of ICAM-1 (green), UEA-1 (endothelial cells, red), and DAPI (blue) staining after 7 days of culture. Co-localization of UEA-1 and ICAM-1 indicates vascular inflammation (yellow). Inflammation of non-endothelial cells is localized by green only staining. Scale bars = 200 μm. **C:** ICAM-1 fluorescence intensity in regions of colocalization with UEA-1 (Vascular ICAM-1). Data is normalized as fold increase over control devices which replace with the TME with a fourth PMN compartment (no cancer in the system). * Indicates p < 0.05. **D:** Normalized vascular ICAM-1 in each compartment of LC-PMN devices. **E:** Relative non-vascular inflammation represented as a ratio of vascular to non-vascular ICAM-1 intensity discriminated based on colocalization with UEA-1. Trend lines in Panels D and E illustrate the linear gradient trends expected in a static culture system.

## DISCUSSION

Fabrication of fluidic culture devices is the primary technical barrier that limits broader adoption of organ chip technologies in preclinical research laboratories. We developed widely adaptable procedures for using low-cost benchtop SLA resin printers to manufacture master molds suitable for PDMS soft lithography. These workflows were developed with the intention of lowering technical barriers to entry in labs without prior microfabrication experience. All organ chip mold designs can be downloaded and used in the designed form or configured according to user specifications (see **Supplemental Information**). Regardless of mold design, the reported post-processing workflow addresses the impact of resin leachates on PDMS curing in resin molds (**Figure 1D**), warping of resin molds and molded PDMS layers (**Figure 1F**), and resin mold surface roughness that dictates optical clarity and hydrophobicity of molded PDMS layers (**Figure 2, Supplemental Figure 2**).

Issues with PDMS curing in SLA printed molds have been well documented across multiple studies utilizing different commercial SLA printers and different resin compositions.^24–26^ While the formulation of proprietary SLA resins are often not published, most commercial SLA resins rely on phosphine-oxide based photo-initiators.^27,28^ Remnants of these photo-initiators may remain unreacted on the surface and within the bulk of SLA prints and interfere with platinum-based silicon curing catalysts.^26^ We found that heating removes the chemicals interfering with PDMS curing and enables flattening of warped molds when coupled with compression by leveraging the malleability of SLA resins that caused warping during the resin curing process (**Figure 1**).

The clarity of fully-cured PDMS microfluidic devices is derived from the smooth surfaces of silicon master molds generated by photolithography in clean room settings.^8,29^ We successfully manufactured devices with near perfect optical clarity using rough surfaced molds refinished with a polyurethane (PU) clear coat (**Figure 2**). Other methods for mitigating 3D printed mold surface roughness include the use of omniphobic lubricants and cycles of PDMS replica molding.^10^ Whilst there are numerous potential approaches smoothen mold surfaces, the use of widely available automotive refinishing products in our workflow ensures that this method is adaptable in any setting where ventilation is controlled. Mold surface roughness also affects liquid interactions in PDMS organ chips, which is of importance for designs that leverage surface tension for membrane-free liquid patterning. Contact angle measurements confirmed increased hydrophobicity of uncoated surfaces, which can be explained by theory that predicts increased liquid repellence which could promote cavitation and liquid spilling over the guide (**Supplemental Figure 2**).^30,31^ Stronger cohesion of the liquid with the PDMS guide structure facilitated by the hydrophilicity of a smoother surface would then be expected to promote pinning of injected liquids along the guide. Whilst PU-coated mold surfaces were clearly smoother upon SEM inspection, molded PDMS from coated and uncoated molds exhibited a microscale surface fracturing pattern that is not typically seen on PDMS surfaces when using standard SU-8 silicon molds (**Supplemental Figure 3**). The PU-coating method corrects optical clarity and enables tuning of surface hydrophobicity, but these surface imperfections of the molded PDMS may impact certain application spaces. Importantly, these PDMS surface features did not negatively impact tissue interface stability or cell performance in MFOC culture.

We established design specifications of geometric features for achieving successful loading of milliscale membrane-free organ chips (MFOC) and determined that bulk tissues can be scaled up to a volume of at least 48 mm^3^ without any change in loading efficiency (**Figure 3**). We did not test larger tissue volumes, but our data suggests that the MFOC design specification of h/H = 0.5 can enable engineering of larger tissue volumes in a fluidically accessible and vascularizable format. The finding that h/H = 0.75 design commonly used in microscale MFOC was intractable for milliscale MFOC loading highlights the impact of total liquid volume and mass on surface tension-guided liquid patterning. We designed milliscale MFOC for the downstream development of vascularized MPS. Microfluidic models with perfusable vasculature have been reported in numerous formats for more than a decade.^17,18,32,33^ Our study expands on this previous work by establishing milliscale models with perfused vasculature spanning more than one millimeter (**Figure 5**). Notably, we successfully achieved patent anastomosis in our system with static culture using standard vascular cell medium formulations (**Figure 5**). There are natural fluctuations of pressure that can drive flow in the anastomosed vessels in static culture conditions, due to slight differences in media reservoir volumes and device tilting during handling for feeding and imaging. The impact of shear stress alterations on endothelial barrier function is a tenet of vascular physiology, and mechanical conditioning of engineered microvessels via flow induced shear stress has been shown increase the maturation of endothelial barriers.^21,34,35^ Our study exploring flow conditioning (**Figure 6**) corroborates previous reports of improved barrier formation with syringe pump-driven flow when compared to the more convenient method of device rocking that produces oscillatory bidirectional flow.^34^ Importantly, those studies were performed using engineered single microvessels, which allow for precise determination of applied shear stresses. Whilst correlation of applied shear stresses with permeability in the 3D vasculature of our milliscale MFOC models is more challenging, our results suggest that continuous unidirectional flow is the optimal method to accelerate and enhance barrier maturation in engineered perfusable vasculature.

Modeling the systemic effects of malignancy is an integrated fashion will eventually require the development of minimally complex body-on-a-chip systems for investigating relevant distant organ effects. Cancer pathophysiology involves systemic processes such cachexia and premetastatic niche cultivation, which are driven in part by inflammation that originates at the site of the primary tumor and propagates to regional and distant sites.^36–38^ We previously established a microphysiological model of tumor cytokine-driven muscle tissue injury, a cardinal feature of cancer cachexia, using a device that fluidically interconnects an engineered tumor microenvironment with patterned muscle tissue constructs.^39^ The LC-PMN model provides a design framework and tractable starting point for developing multi-organ models with an integrated vasculature (**Figure 7**). It is important to note that the LC-PMN model was operated in static culture. Future work will test pumping and rocking culture regimens to tune mass transport in the system.

Limitations of the current manufacturing approach are primarily related to attainable printer resolution and potential material differences with resins used in other printers. The post-processing workflow we report is broadly applicable to any materials which may contain trapped volatiles or are prone to distortion. MFOC in various forms will continue to evolve as a platform technology for creating MPS with the critical physiological component of a perfusable vasculature. The systems and methods we report will allow non-expert laboratories with access to inexpensive SLA printers and basic fabrication supplies to design, fabricate, and test the usability of MFOC for specific disease modeling and drug delivery applications. The impact of externally applied flow conditions on barrier function requires more systematic investigation to establish precise engineering control, although it is a challenging modeling problem from an experimental and theoretical viewpoint due to the heterogeneous 3D architecture of bulk vasculature and the resultant fluctuations of local flowrates and shear stresses within the tissue. The results attained using HUVEC and HLF as the stromal-vascular cell source need to be replicated using organ-specific combinations of fibroblasts and endothelial cells to create organotypic models. Future versions of the LC-PMN model will replace lung carcinoma cell spheroids with LC patient biopsy derivatives. Our long-term goal is to integrate engineered organ compartments such as muscle, liver, and adipose tissue in the prototype LC-PMN model on the path toward developing a minimally sufficient body-on-a-chip system with integrated vasculature for modeling the systemic effects of malignancy and screening anti-metastasis and anti-cachexia therapies.

## Materials and Methods

### SLA Printing

Device molds were drafted as 3D drawings using SolidWorks (Dassault Systèmes) or Fusion 360 (Autodesk). Molds were created in a top-down view and printed using a Form 3B SLA Printer (Formlabs). Final designs were exported at “.STL” files to Formlab’s PreForm software. Formlab’s PreForm software was used to prepare 3D drawing files for printing. Each part file was oriented so that the mold base was parallel with the printer’s build platform. Print supports were autogenerated within PreForm with a 0.65 mm touchpoint size and a 1.30 support density. All molds were printed using Formlab’s proprietary ‘Grey’ or ‘Clear’ SLA resins. CAD files for all organ chip mold designs used in these studies are available in downloadable form along with a list of dimensions for all designs in the **Supplemental Information**.

### Mold Post-Processing

Completed prints were washed in isopropyl alcohol (IPA) according to Formlabs protocols for the resins used, and then washed in IPA for 20 additional minutes in the FormWash (Formlabs). Post wash, molds were dried until IPA was completely evaporated. Molds were then dried and cured under UV light (FormCure, Formlabs) at 60°C. Clear resin parts were cured for 15 minutes. Grey resin parts were cured for 30 minutes. Warping caused by distortion of the printing process and the curing process was corrected by first baking Clear resin molds at 130°C for 2 hours and baking Grey resin molds for 3 hours. For the last 30 minutes of bake time, 2 stainless steel jeweler’s blocks were added to the oven to heat. Molds were removed and placed between two jewelers’ blocks with or without clamping. Flatness of the parts was assessed visually relative to a straight edge. We used a commercial painting airbrush (Model 105 Patriot Fine Gravity Airbrush, Badger Airbrush Co.) to coat molds with lacquer thinner (Klean-Strip). Parts were dried for 15 minutes while preparing automotive clear coat (Finish 1 FC720/FH612, Sherman Williams) mixed at a 1:4 ratio of hardener to clear coat. The mixture was airbrushed from approximately 8 inches away using a continuous back and forth motion and then a continuous up and down motion until a thin layer of clear coat was visible. Four layers were applied with the mold rotated 90° between applications. Coated molds were dried for 6 hours before silanization according to standard protocols for PDMS soft lithography. ^40^

### PDMS soft lithography

PDMS (Sylgard 184, Ellsworth Chemical) was mixed at a 1:10 ratio of PDMS curing agent to PDMS elastomer by weight and degassed. PDMS soft lithography was carried out according to standard protocols used for silicon master molds.^9^ To produce PDMS molds with two flat surfaces, we applied a cleaned 2×3 inch glass slide to the top of the mold to sandwich the uncured PDMS, carefully avoiding bubble formation. A jeweler’s block was placed on top of the slide to ensure a tight seal and filled molds were baked at 60°C for at least 4 hours.

### Device assembly

Cured PDMS device layers were removed from molds and detached from the glass slides using a razor blade. Clear packing tape was used to remove any dust from the PDMS parts. Through features, such as inlets, were removed using biopsy punches and an X-acto knife as needed. Device bonding was achieved by PDMS stamping as reported previously. ^41^ Briefly, 3 g of 1:10 PDMS was spin coated on a petri dish to create a thin stamping layer. Device layers were manually stamped onto this PDMS layer and then stacked together. Reproducibility of layer alignment was ensured by the inclusion of Lego-like press fitting features (**Supplemental Video 2**). Assembled devices were cured at room temperature for at least 24 hours.

### Fabrication of Double Guide MFOC

The upper guide structure in the double guide MFOC design required generation of a second mold. Due to the negative feature resolution limitations of our SLA printer, we bisected the symmetrical design and printed two molds that were mirror images of each other. PDMS parts cured in these molds were PDMS stamped and sandwiched together with the guide structures facing inward. Cap layers containing media reservoirs were fabricated and PDMS stamped to the top of double guide MFOC to complete the final assembly prior to use in culture. Alternatively, PDMS replica molding of positive feature prints can be used to fabricate the upper guide rail layer (see **Supplementary Information**).

### Injection testing

To test the usability of each design iteration, the devices underwent a series of injection testing. In each trial, at least 6 users with varying levels of familiarity with injecting microfluidic devices were recruited. Three replicates of each design were fabricated. We used 200 μl pipettes with standard tips to inject food dye solutions. Following injection, the water was removed from the device, and the device was completely dried using compressed air. If injected water reached the outlet without breaching into side channels, the trial was successful. Each user completed 10 tests per design and loading success rates were recorded. A 100 µg/mL solution of 40 kDa FITC-dextran (Sigma-Aldrich) was mixed into a collagen hydrogel prepared as described above and injected into the central channel. We acquired 3D LSCM stacks to visualize tissue interfaces on the guide structures.

### Cell culture

Human umbilical vein endothelial cells (HUVEC) (ATCC) were maintained in Vascular Cell Basal Medium supplemented with Endothelial Cell Growth Kit-VEGF (ATCC). Human lung fibroblasts (HLF) were maintained in Fibroblast Basal Medium supplemented with fibroblast Growth Kit-Low Serum (ATCC). A549 lung adenocarcinoma cells were cultured in F12 medium with 10% FBS. All culture media contained 1% antibiotic-antimycotic (Corning). Cells were maintained in a humidified tissue culture incubator maintained at 37°C and 5% CO_2_ and used at passages 3-7.

### Organ Chip Seeding

Organ chips were exposed to UV light in a cell culture hood for 1 hour. The surfaces of chambers of multilayer devices and lanes of MFOC that house bulk 3D tissues were functionalized for ECM hydrogel anchorage using the polydopamine (PDA) coating method as previously described.^42^ Briefly, dopamine solution was prepared by mixing 10 mg of dopamine hydrochloride (Sigma-Aldrich) -with 10 mM Tris-hydrochloride (Sigma-Aldrich). Dopamine solution was sterile filtered and injected in the tissue chambers and lanes of devices. Devices were incubated for 2 hours at room temperature in the dark, aspirated and used immediately. Multilayer membrane-bound organ chips were loaded with collagen type I hydrogel (2.5 mg/ml) containing primary normal human lung fibroblasts (5×10^5^ cells/ml) in the 3D tissue chamber, and the upper tissue layer channel was seeded with A549 lung adenocarcinoma cells (1 × 10^6^ cells/ml). MFOC were seeded with HLF and HUVEC (2×10^6^ cells/mL each) admixed in 2.5 mg/mL collagen I (Corning), 5 mg/mL fibrinogen (Sigma-Aldrich), and 1 U/mL thrombin (Sigma-Aldrich). This cell inoculated hydrogel precursor was injected into central tissue lane of MFOC devices and incubated for 30 minutes prior to filling the side channels and media reservoirs with endothelial cell growth medium (VEGF kit) containing 25 µg/mL aprotinin (Sigma-Aldrich) to inhibit fibrin degradation.

### Establishing Anastomosed and Perfusable Bulk Vasculature

After two days of bulk tissue vasculogenesis initiated as described above, HUVEC (10×10^6^ cells/mL) were injected into the MFOC side channels. Devices were maintained in static culture for another 7 days with media changes ever 24 hours to allow for lumen formation and vessel maturation. We define static culture as no pumping or rocking but slight differences in liquid head height of medium reservoirs lead to trickling flows in the anastomosing vasculature. One channel was loaded with 100 µg/mL of 40 kDa FITC-dextran (Sigma-Aldrich) in PBS with the reservoirs filled to an elevated pressure head and allowed to perfuse through vasculature until it reached equilibrium with the opposite channel. Videos were acquired as time series of the DIC and FITC channels using the Nikon C2 LSCM. 3D stacks were acquired to visualize patency of the vasculature throughout the 3D bulk tissue.

### Fluid Flow Conditioning of Anastomosed Vasculature

We developed an alternative protocol of seeding the side channels as described above on Day 0, immediately after polymerizing the bulk tissue. After allowing cell adhesion and integration overnight, flow is applied across the bulk tissue using a laboratory rocker or syringe pump for 4 days. A laboratory rocker was placed in an incubator and set for a 15-degree incline at 1 cycle/min. Devices are oriented with central channel in line with the axis of rotation. For pump-driven conditioning, a syringe pump was set to pull 20 ul/hr from one side of a device. The opposite channel of the device was connected to a liquid media reservoir. Tygon tubing and stainless-steel fittings are used to connect devices to the syringe pump and reservoir.

### Perfusion and Permeability Quantification

Perfusion throughout vascular networks is assessed using a FITC-dextran solution and confocal microscopy. A solution of 40kd FITC-dextran is diluted 1:250 into media is first prepared. One device is mounted on the stage of a confocal microscope. Liquid media channels and reservoirs are emptied, then one channel is filled with FITC-dextran solution; the reservoir is filled to maintain consistent hydrostatic pressure across each trial. DIC-confocal microscopy is used to capture a 10-minute time series at 10x magnification at 1 frame/5s. Large composite images are taken after each time series. To quantify perfusion and diffusion throughout the vascular network, FITC fluorescence is measured in the intravascular and extravascular space. A DIC image at t=0 is used to create a binary mask of the vascular area. FITC fluorescence within the masked area is considered intravascular and all other FITC fluorescence is considered extravascular. All computation were performed in MATLab. The alkaline lignin-g-PLGA nanoparticles (100-200 nm size range) entrapped with Cy5 used for perfusion testing were provided as a gift by Dr. Cristina Sabliov.^43^ Briefly, particles were washed, resuspended in media at 1 mg/mL, and sonicated for dispersion. Nanoparticle solution was then injected into the anastomosed vasculature using the same method for FITC-dextran perfusions described above and LCSM stacks were acquired for visualization.

### Whole Mount Staining in Organ Chips

Tissue layers and bulk tissues in organ chips were stained using adaptations of previously reported protocols.^39,44,45^ Live tissues were incubated with 4 μM calcein-AM for 30 minutes (Live/Dead, Invitrogen) to visualize viable cells. For all other stains, tissues were fixed by loading 4% paraformaldehyde in the organ chip liquid channels and incubating for 1 hour at room temperature then overnight at 4 °C. Devices were washed with PBS and stored at 4 °C prior to staining. For convenient visualization of the vasculature, tissues were stained with a cocktail of 4 μL/mL 4′,6-diamidino-2-phenylindole (DAPI) to label nuclei, 4 μL/mL Alexa488-conjugated phalloidins to label actin in all cells, and 20 μL/mL DyLight594-conjugated Ulex Europeas agglutinin I (UEA-1, Vector Laboratories) to specifically label endothelial cells.^46^ The staining cocktail was prepared in 1X PBS with 0.2 % Triton-X and 1% BSA. Devices were loaded with the staining cocktail, rocked gently for 1 hour at room temperature, refrigerated overnight, and rocked for an additional hour at room temperature before washing. Whole mount Immunohistochemistry for ICAM-1, laminin, and Ki67 was performed by adding a block and permeabilization step with 3% BSA and 0.2% Triton-X in 1X PBS for 30 minutes. Rabbit anti-Ki67 (Abcam, ab15580), Mouse anti-ICAM-1 (Abcam, ab171123), or Rabbit anti-laminin (Abcam, ab11575) were diluted 1:100 in 1% BSA with 0.1% Triton-X. Primary antibodies were gently rocked for at least 4 hours before overnight incubation at 4 °C. After washing with multiple changes for 2-3 hours, anti-Rabbit or anti-Mouse secondary antibodies (Abcam) diluted 1:250 in the same antibody buffer were incubated for 2 hours with optional phalloidin and DAPI counterstaining as described above. After several changes of wash buffer samples were briefly checked for removal of background fluorescence and stored loaded with PBS in foil wrapped dishes at 4 °C until imaging.

### Imaging and Image Analysis

Stained tissues were fixed in position on a slide and imaged on an inverted Nikon C2 laser scanning confocal microscope (LSCM) equipped with a Nikon DS-FI3 camera. All images for fluorescence intensity quantification were collected at fixed exposure times and laser intensities. Max intensity projection Z-stacks were exported as TIFF files with no LUTs attached. All Image analysis was completed in MATLAB (R2021b). All morphometric image analysis of vascular networks and perfused vasculature was completed in MATLAB (R2021b). We used a pretrained deep neural network to denoise each image and adaptive histogram equalization was used to standardize contrast across the image set.^47^ We used a circular Hugh transform to identify rounded cells in each image and counted rounded cells to assess network completeness.^43^ Rounded cells were removed from images prior to segmentation and morphometric analysis. We smoothed images using an edge preserving filter with a Gaussian kernel and applied a threshold to remove remaining low-intensity noise.^44^ We then segmented pre-processed images and quantified morphometric parameters using an open-source automated segmentation tool.^45^ All devices stained with ICAM-1 were imaged at a fixed laser intensity, and mean pixel intensity was computed for each image. ICAM-1 localization to endothelial cells was discriminated by colocalization with UEA-1 lectin. Mean fluorescence intensity was calculated for the total image, UEA-1 co-localized pixels (vascular inflammation), and pixels with ICAM-1 signal only (non-vascular inflammation).

### Statistical Analysis

All vasculogenesis experiments were repeated n=3 times. A large, composite 1×6 stitched image was acquired from each sample to ensure convergence of morphological variables. Images were preprocessed and segmented per above image analysis methodology. All statistical analyses were completed in GraphPad Prism V 9.2. Comparison of MFOC injection loading success rates, comparison of morphometric parameters between days of vascular network formation, and ICAM-1 fluorescence intensity in PMN model were made using unpaired t-tests with a 95% confidence limit for statistical significance, i.e. p < 0.05. Sample sizes and the number of independent experiments performed are indicated in the respective figure captions.

## Acknowledgements

We acknowledge Max Lobovsky, CEO of Formlabs, for generous provision of technical support for the F3 SLA printer and loaning of a second unit while we optimized printer performance. We also thank Formlabs technical staff for performing test prints with our CAD files at the Formlabs facility and providing useful technical feedback. We thank Dr. Cristina Sabliov in the Department of Biological and Agricultural Engineering at Louisiana State University for the nanoparticles used in perfusion studies.

## Conflicts of Interest

The authors have no conflicts of interest to report.

## Data Availability Statement

All CAD design files, and the code used for image analyses are made available in the Supplementary Information for this article. All data and more detailed explanations of laboratory procedures will be made available upon reasonable request to the corresponding author.

## Supplemental Figures

**Figure S1.**
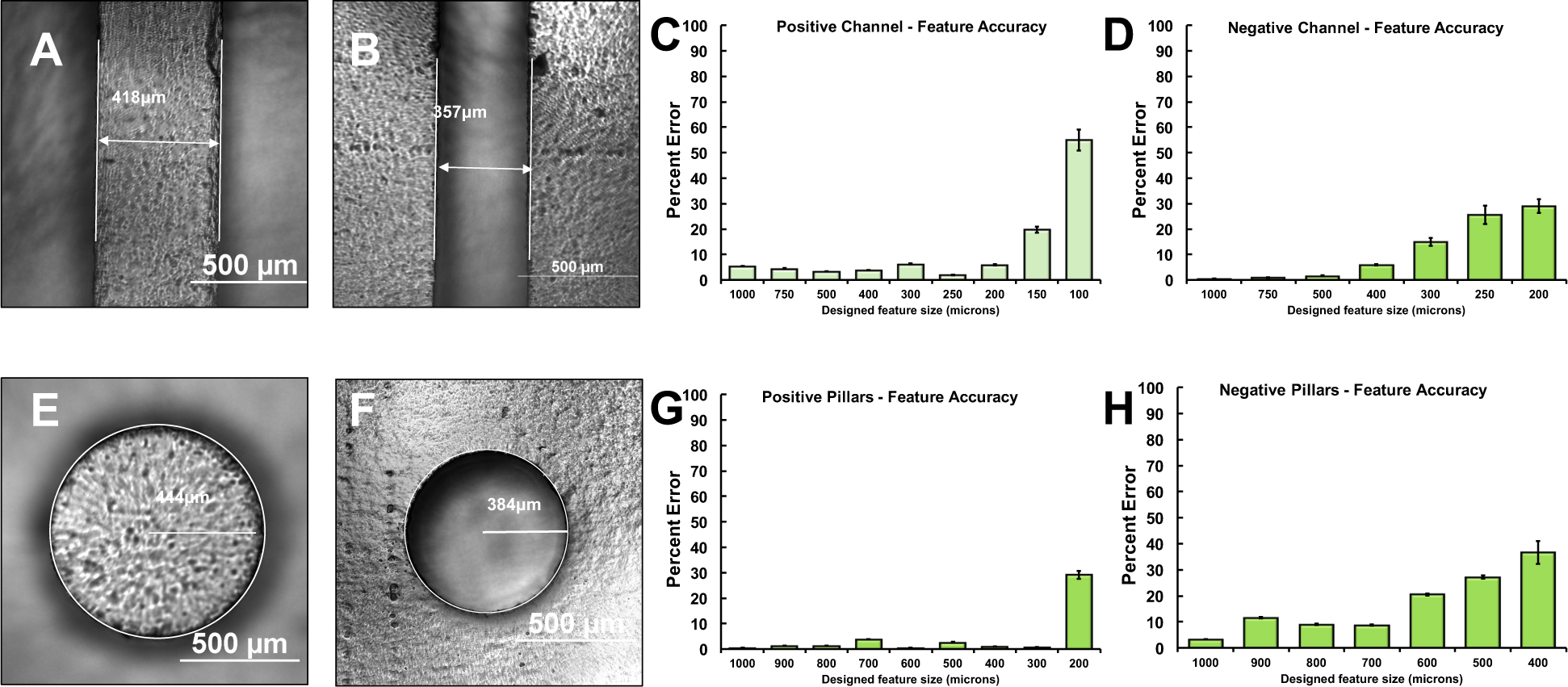
– Formlabs F3 printer resolution testing (supplement to Figure 1. We used a Formlabs F3 SLA printer with Grey and Clear resins as described in **Materials and Methods**. We first determined the build resolution of the printer with these resins focusing on features commonly used in microfluidic device molds such as rectangular channel features and cylindrical features used to mold fluidic access ports. We tested the printer resolution for both positive features built above the floor of the mold and negative features fabricated as hollow impressions below the floor of the mold. Please note that these data reflect usage of the printer in our hands with the specific designs tested and do not represent the absolute limits of the printer using these materials. We measured the built feature dimensions using a stereomicroscope and calculated a percent error relative to the designed dimensions (**Figure S1A, B, E, F**). Less than 10% error was designated as acceptable accuracy for organ chip mold fabrication. We found that the positive feature resolution for both rectangular channels and cylindrical pillars was accurate down to sizes of 200 µm (**Figure S1C, G**). These results confirmed that the printer has excellent resolution for our intended applications of manufacturing molds for millifluidic organ chip devices. Negative channel features were accurate down to 400 µm, whereas negative pillar features were inaccurate at sizes as large as 900 µm (**Figure S1D, H**). These data indicate that negative feature resolution is significantly impaired relative to positive feature resolution, thereby necessitating designs that avoided negative features or implementation of a PDMS replica molding technique that enables fabrication of negative feature PDMS molds with resolution that matches the positive feature build accuracy of our SLA printer (**Figure S2**).

**Figure S2.**
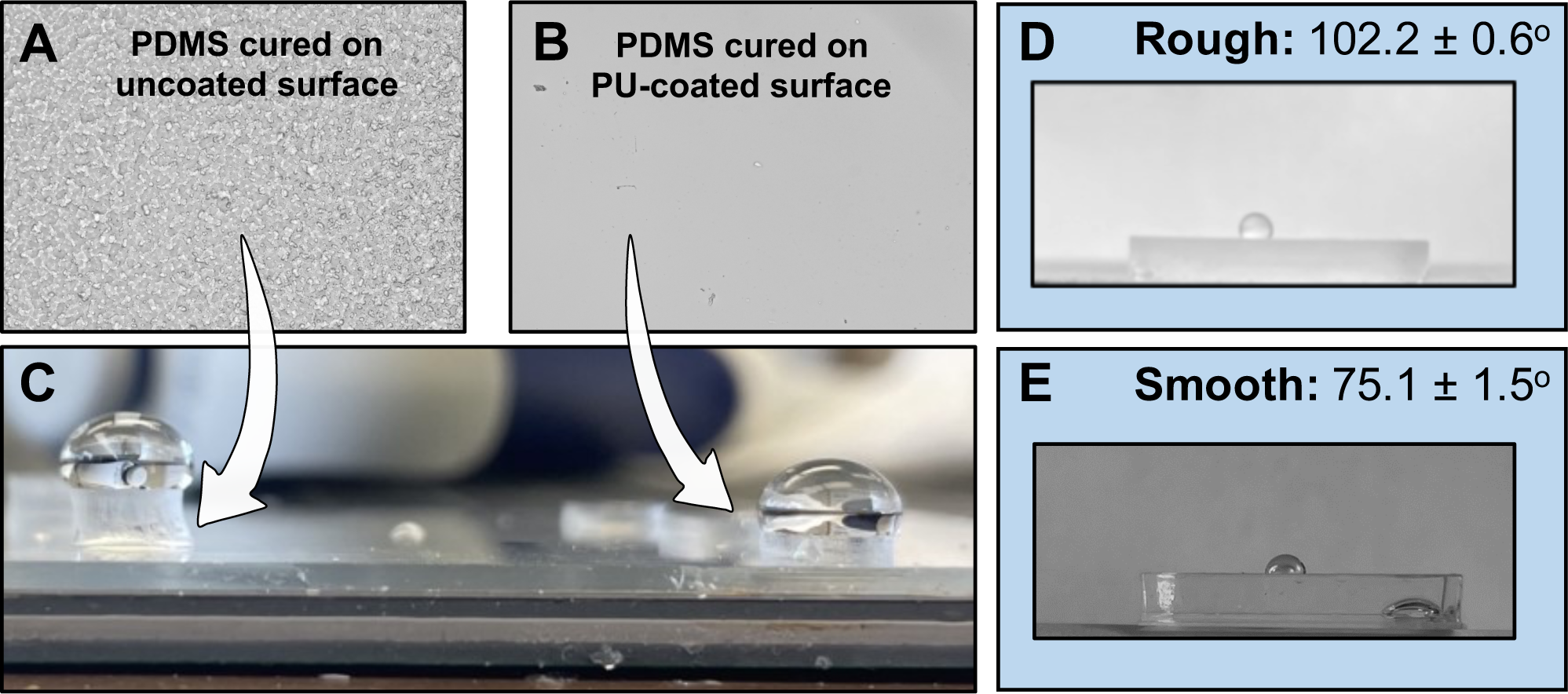
– Contact angle of PDMS from uncoated and PU-coated molds. We compared the surface wettability of PDMS cured on uncoated mold surfaces and PU-coated mold surfaces to better understand the result of increased loading failure rates in MFOC fabricated using uncoated molds (**Figure 2F**). Differential interference contrast imaging confirmed the roughness of PDMS surfaces from uncoated molds (**Figures S3A, B**), which resulted in a visually increased hydrophobicity compared to PDMS surfaces from PU-coated molds (**Figure S3C**). Increased hydrophobicity of the rough PDMS surfaces from uncoated molds was confirmed by contact angle measurements (3 replicate surfaces, 10 drops per surface) (**Figures S3D, E**). The low variance in the contact angle data confirms the consistency of surface properties of SLA-printed molds and the PU coating procedure. Statistical analysis revealed that PDMS surfaces from PU-coated molds are significantly more hydrophilic (75.1 +/− 1.5° vs. 102.2 +/− 0.6°, P < 0.001).

**Figure S3.**
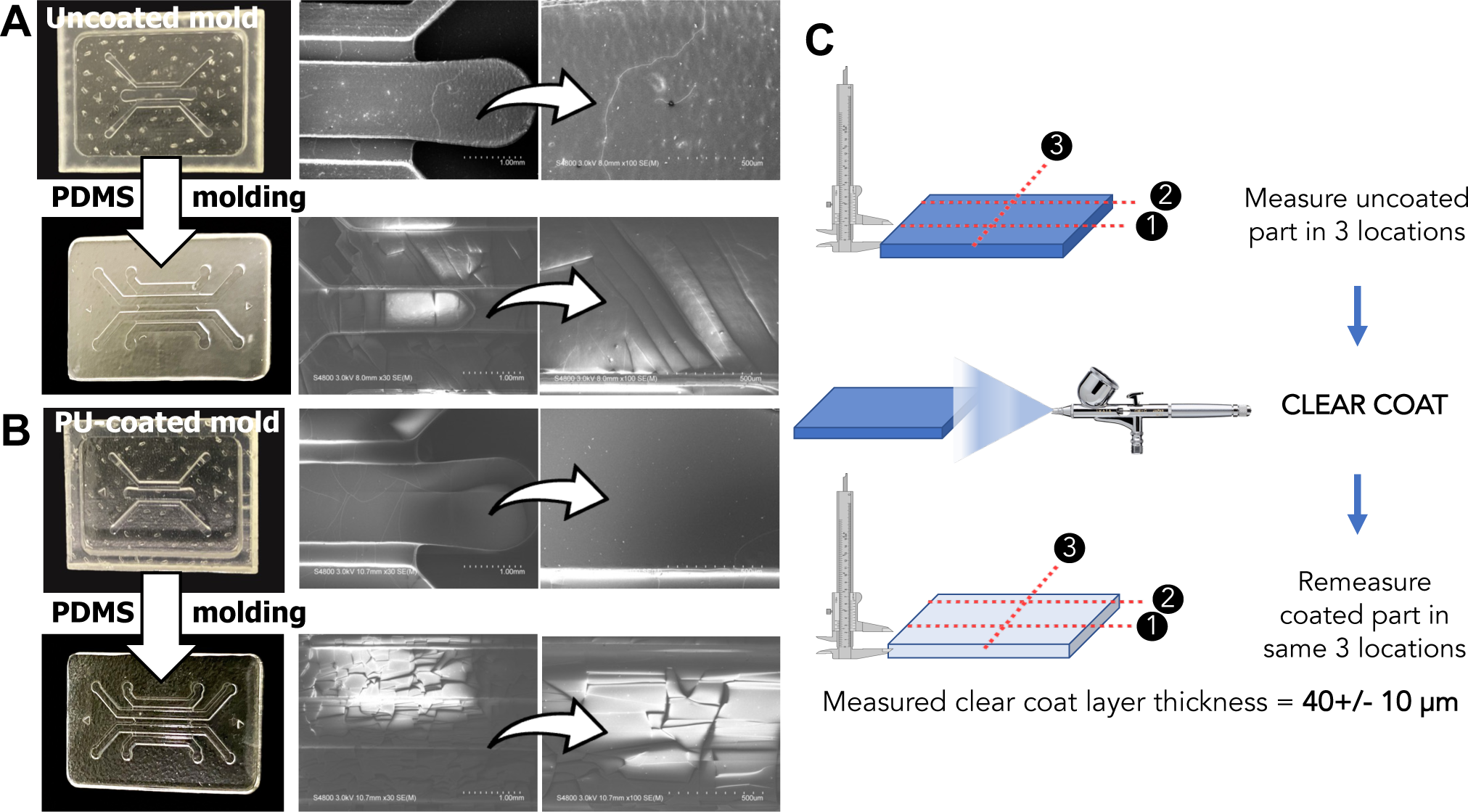
– SEM imaging of uncoated and polyurethane (PU) coated SLA resin molds and the resultant molded PDMS surfaces, and measurement of PU coating thickness (supplement to **Figure 2**. **A:** Uncoated ‘Clear’ resin mold and a molded PDMS device layer after soft lithography using the same mold. The opacity of the molded PDMS layer is grossly observable (left, digital photographs). Scanning electron microscopy (SEM) imaging revealed rough texture (top right). SEM revealed that the surface of molded PDMS from uncoated molds has a fractured appearance with visible fissures (bottom right). **B:** PU coated ‘Clear’ resin mold and a molded PDMS device layer after soft lithography using the same mold. The clarity of the molded PDMS layer is grossly observable (left, digital photographs). SEM revealed a markedly smoother surface compared to PDMS layers from uncoated molds (top right panels). The same fissured surface was seen on SEM of PDMS layers from PU coated molds, despite the functional optical clarity (bottom right panels). This observation indicates that these fissures are not the direct cause of the opacity of PDMS from uncoated molds. **C:** A micrometer was used to measure thickness before and after PU coating. Thickness was measured at 3 locations. PU coating thickness (see **Methods** for details of coating procedure) was estimated to be 40 +/− 10 micrometers (n = 3 separate coated surfaces).

**Figure S4.**
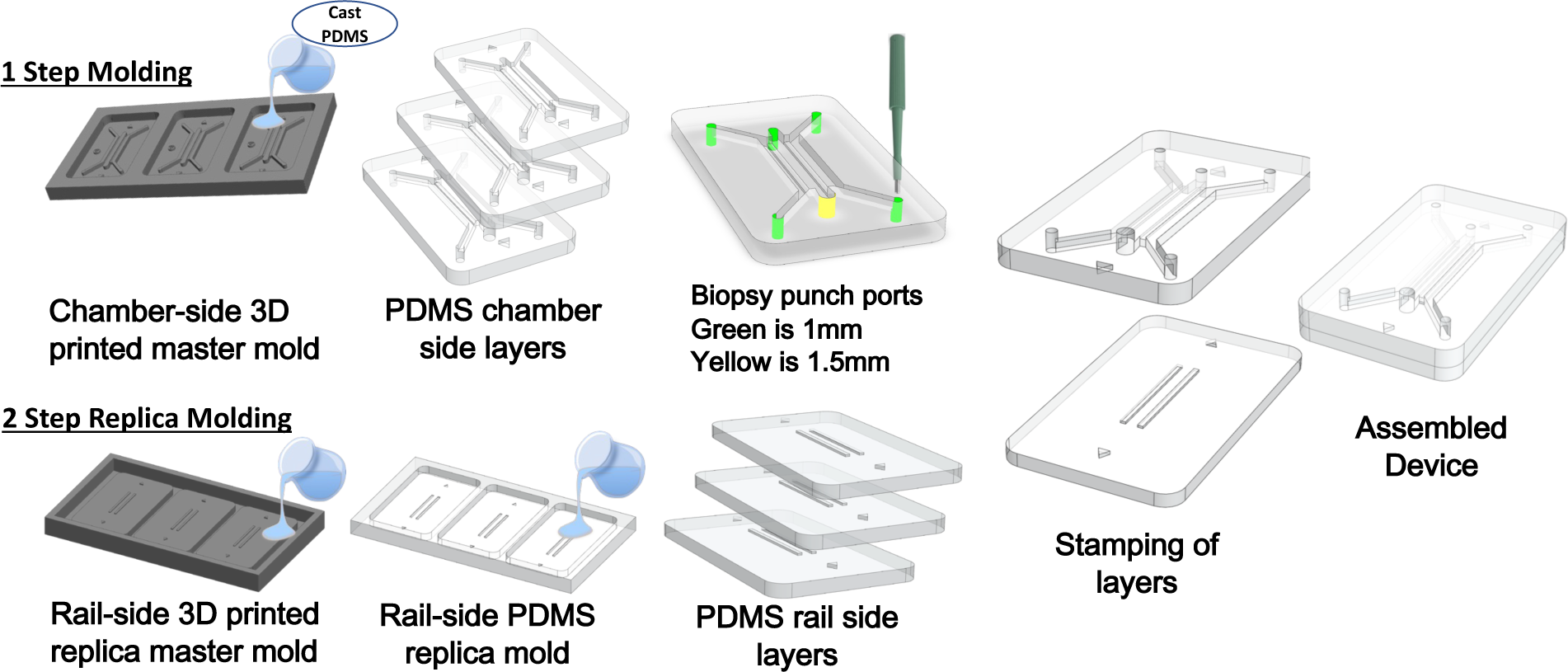
– Creating PDMS replica molds to achieve negative feature molding that matches the F3 SLA printer’s positive feature build accuracy (supplement to **Figure 2, 3**) PDMS replica molding was performed as described in **Supplemental Methods**.

## Supplemental Videos

**Supplemental Video 1** – Rotating view of 3D tissue interface assembly in the organ interstitium chip seeded with human lung fibroblasts in collagen type I hydrogel in the interstitial layer and lung adenocarcinoma cells in the upper channel (supplement to **Figure 1**). Video render of a confocal z-stack spanning 1 millimeter in the z-direction. Actin labeling with Alexafluor-488 phalloidins illuminates the cytoskeleton of all cells. Regions of slices from this stack are shown in **Figure 1H**.

**Supplemental Video 2** – Reproducible assembly of MFOC without the use of imaging guidance (supplement to **Figure 2**). We included triangular Lego-like features that enable effortless and reproducible press fit bonding with accurate alignment of features in bonded layers of MFOC devices, which is strictly required for proper surface tension patterning. MFOC are stamped with a thin coating of uncured PDMS and press fit as shown in the video. No additional steps are needed to facilitate or confirm layer alignment.

**Supplemental Video 3** – MFOC loading to demonstrate device functionality and exemplary testing results (supplement to **Figure 2** and **Figure 3**). The first clip shows a double guide MFOC with h/H = 0.5, H = 1 mm, and W = 1.2 mm being loaded with green food dye in water. The device was loaded by gravity driven flow with a pipette tip filled to 200 µl to create a large pressure drop beyond what occurs during careful manual pipette injection. The second clip shows a successful manual pipette loading during user testing. The third clip shows a failed manual pipette loading during user testing.

**Supplemental Video 4** – FITC-dextran perfusion of anastomosed vasculature in MFOC (supplement to **Figure 7**). Tissues with anastomosed vasculature were produced as described in **Methods** and shown in **Figure 7**. This video shows a time lapse of 15 minutes of perfusion after introducing 20 kDa dextran into one of the MFOC side channels. The absence of dextran leakage demonstrates patency of the anastomosed vasculature formed in a fully static culture system. Flow conditioning of the network using rockers and pumps can influence the rate and degree of endothelial barrier formation (1,2).

## Organ chip design specifications

This section contains design specifications and links for downloadable STL files. All devices are fabricated as described in Materials and Methods. Briefly, the multilayer organ chip with polyester membranes is vertically layered and requires the placement of manually trimmed porous membranes (such as track-etched polyester with 0.4 µm pores) that cover channel and tissue chamber features. All devices are bonded by PDMS stamping.

Key dimensions of Organ Chip designs are available in the downloadable folder of STL files.

*Link to be added upon acceptance for publication. Email corresponding author in the interim.

## Supplemental Methods

### Scanning Electron Microscopy

Samples were coated with Cressington 208 Carbon Coater and imaged with Hitachi S-4800 Field Emission Scanning Electron Microscope (FESEM, S4800 Hitachi, Japan) at 3 keV. The surface topography of PU-coated versus non-PU-coated molds and the corresponding molded PDMS device layers were observed and analyzed using a secondary electron beam at 10 uA at x30 and x100 magnification.

### PDMS replica molding

Double guide MFOC designs used for these studies required a second mold generated using PDMS replica molding, due to negative feature resolution limitations of our SLA printer. Instead of the 3D printed mold being the complement of the final PDMS part, the mold was a replica of the desired PDMS part (**Figure S7**). The PDMS replica mold of the eventual MFOC device layer was created using the PDMS soft lithography methods described above. The PDMS replica mold part was silanized using standard methods prior to replica molding the device layers.

## MATLAB code for image analysis

*Link with all scripts and sample image sets for testing will be added upon acceptance for publication or upon reviewer request.

These scripts were written for image preprocessing, counting of non-networked cells, and plotting of morphological parameters.

Built in MATLab R2020b (Windows) with deep learning toolbox.

Instructions for Use --------------------------------

1. RUN VascProgPreprocess
2. RN Roundcell_ID
3. Segment Images in ’…\ MC_VascProg_ImageProcessing\CIRCLE_id\network_extract’ FOLDER-REAVER was used for segmentation, quantify and export morphological metrics
4. RUN VascProg_DataAnlysis to produce matlab plots of REAVER metrics

Matlab Script Descriptions --------------------------------

…

VascProg_DataAnalysis ------

-- denoises images with CNN, applies adaptive histogram equalizaiton for contrast adjustment, smooths and filters noise

- this function prepares raw confocal images for segmentation using REAVER

- parameters may be adjusted to accomodate specific image aquisition settings; image aquisition and staining protocol should be consistent across entire dataset

Roundcell_ID --------------

-- identifies round cells using a Hugh transform

- this cell counting method standardizes cell counting in an image with diverse cell morphology

- removes round cells from each image and saves resulting image (network only) to ’…\ MC_VascProg_ImageProcessing\CIRCLE_id\network_extract’

- parameters (max/min radius, sensitivity) can be adjusted to optimize cell count accuracy OUTPUTS: ’im_meta’ - metadata for cell count; includes file name, extension, object (cell) location, corresponding cell radius, cell count, approximate cell density [cells/mL]

VascProg_DataAnalysis -----

-- produces matlab plots for REAVER morphological metrics

## Notes

### Competing Interest Statement

The authors have declared no competing interest.

